# Latent representation of the human pan-celltype epigenome through a deep recurrent neural network

**DOI:** 10.1101/2021.03.08.434446

**Authors:** Kevin B. Dsouza, Adam Y. Li, Vijay K. Bhargava, Maxwell W. Libbrecht

**Affiliations:** Department of Electrical and Computer Engineering at the University of British Columbia; Department of Bioinformatics and Systems Biology at the University of Amsterdam; School of Computing Science at the Simon Fraser University

**Keywords:** Epigenomics, Deep recurrent neural network, Genome annotation, Unsupervised learning

## Abstract

The availability of thousands of assays of epigenetic activity necessitates compressed representations of these data sets that summarize the epigenetic landscape of the genome. Until recently, most such representations were celltype specific, applying to a single tissue or cell state. Recently, neural networks have made it possible to summarize data across tissues to produce a pan-celltype representation. In this work, we propose Epi-LSTM, a deep long short-term memory (LSTM) recurrent neural network autoencoder to capture the long-term dependencies in the epigenomic data. The latent representations from Epi-LSTM capture a variety of genomic phenomena, including gene-expression, promoter-enhancer interactions, replication timing, frequently interacting regions and evolutionary conservation. These representations outperform existing methods in a majority of cell-types, while yielding smoother representations along the genomic axis due to their sequential nature.

## 1 Introduction

SEQUENCING-based assays such as ChIP-seq, ATAC-seq, and DNase-seq have recently been used to characterize the epigenome of hundreds of human cell types. These assays detailing a varied number of epigenomic functions like methylation status, local chromatin accessibility, histone modifications, factor binding and chromatin structure are hosted by consortia such as Roadmap Epigenomics [1] and ENCODE [2]. These data sets necessitate integrative methods that summarize them into a useful representation. A popular existing type of method is segmentation and genome annotation (SAGA) algorithms such as Segway [3], ChromHMM [4] and others [5], [6], [7], [8], [9], [10], which produce an annotation of the epigenome of a given cell type.

Existing SAGA annotations are celltype-specific; that is, they annotate activity in a given cell type. This corresponds poorly to most conceptualizations of genomic elements. Other genome annotations, such as annotations of coding genes, are pan-celltype, and it is common to say that a locus “is an enhancer”. Moreover, connecting a genetic locus to a phenotype or disease requires an understanding of its function across different cell types. Existing SAGA algorithms cannot be adapted for this task because they use simple discrete or linear models that cannot capture the complexity of the epigenome across all cell types.

We propose a method called Epi-LSTM that produces a pan-celltype low-dimensional representation of the epigenome. This representation assigns a vector of features to each genomic position that represents that position’s activity across all tissues. This representation encapsulates all information about epigenomic activity across all cell types. Therefore, any consequence of epigenetic activity can be extracted from the representation, including identifying regulatory elements or connecting disease-associated variants to causal functional elements.

We do this using a deep long short-term memory (LSTM) [11] recurrent neural network autoencoder to reduce all existing epigenome data into a single low-dimensional representation. Epi-LSTM uses an autoencoder architecture in which aims to produce a representation that can be used reconstruct the original data as accurately as possible.

There exists a feed-forward autoencoder model called GSAE (gene superset autoencoder) [12], but GSAE was designed with the goal of discriminating tumor sub-types by interpreting gene expression data, taking inter-gene sets association into consideration and doesn’t work with epigenomic data. One neural network representation learning method for epigenomic data exists, called Avocado [13]. Like our method, Avocado produces a representation of the epigenome that assigns a low-dimensional vector to each genomic position. Avocado was initially developed for imputation. It uses distinct embeddings for each cell type, assay type and genomic position, and couples this with a feed-forward neural network that imputes unperformed assays. REFINED (REpresentation of Features as Images with NEighborhood Dependencies) [14], is a method for representing high-dimensional feature vectors as 2D images that can be processed by Convolutional Neural Network (CNN)-based pipelines. Although REFINED is not directly comparable to Epi-LSTM, we use it as a preprocessing step treating the cell-type axis as it’s primary input, the output of which is fed to a CNN autoencoder [15]. We call this REFINED+CNN.

Epi-LSTM’s use of a sequential model brings a number of benefits relative to these methods. The model captures the the spatial relationship of neighboring genomic positions and bypasses the need for fixed input windows. This results in smoother representations along the genomic axis when compared to other methods (Results), which aid in use and interpretation. As a result, the output representations outperform those from other methods at classifying genomic phenomena in a majority of cell-types (Results). In addition, because the model uses an autoencoding backbone, it can be applied to genomic positions that were not used in training by inputting the relevant data into our encoder.

We demonstrate the utility of the Epi-LSTM representation through several analyses. First, we show that this representation simultaneously captures cell type-specific activity across many cell types, including gene expression, replication timing and chromatin contacts. We do this by demonstrating that all of the above phenomena are accurately predictable using just the latent representation. Second, we show that this latent representation distinguishes functional and non-functional regions by showing that the representation accurately identifies conserved regions. Third, we demonstrate how a sequential model leads to smoother and more interpretable representations than existing methods, which do not capture dependence among neighboring positions. A graphical abstract (Figure 1) explains the overall idea behind the work. The genomic datasets which are stacked, are fed into the LSTM Autoencoder framework, the latent representation from which is used for varying downstream genomic tasks. The framework is further elaborated in section 3.2.

**Fig. 1:**
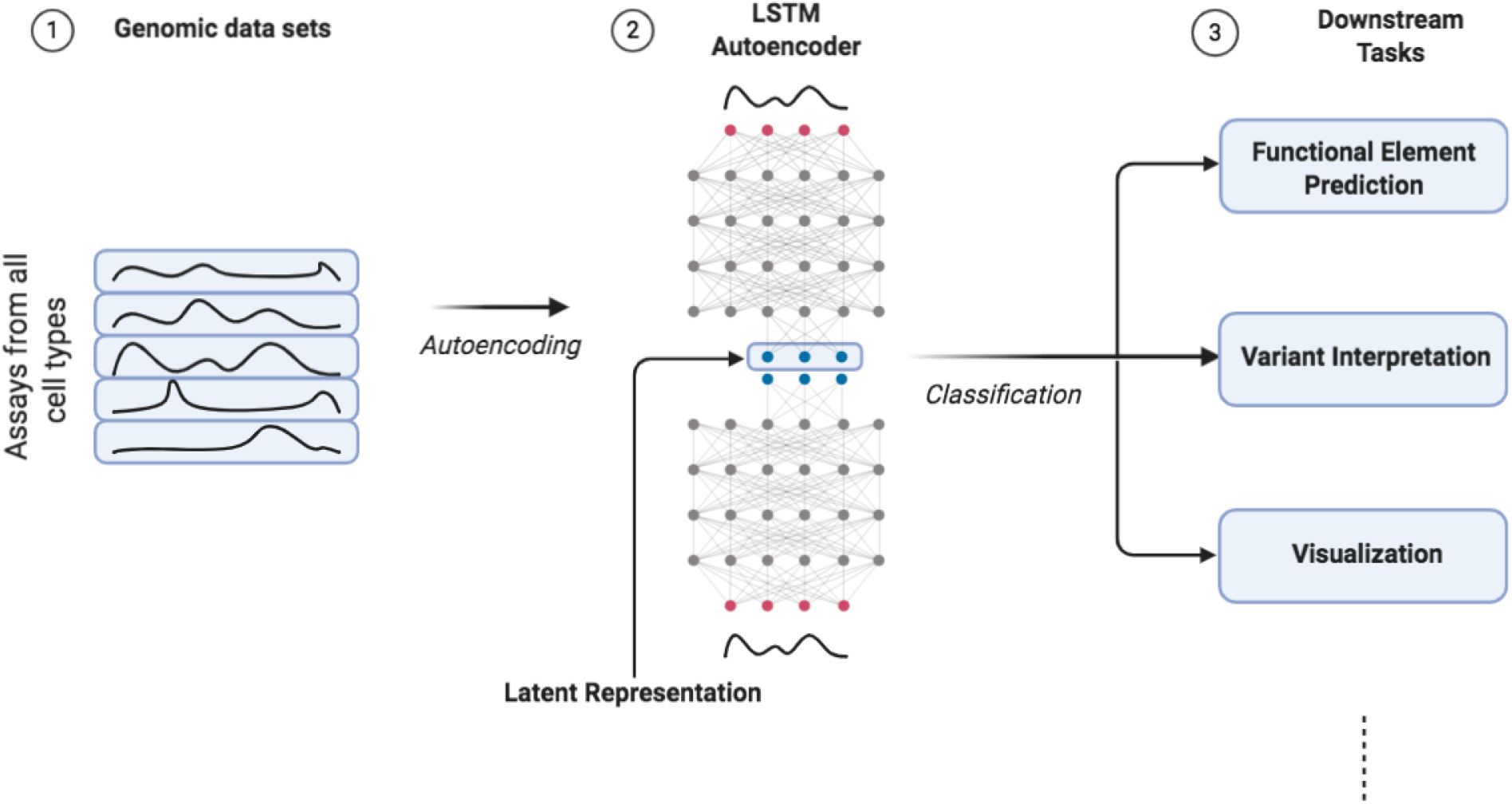
The overall idea behind the work. The epigenomic datasets from all cell types are fed into an LSTM autoencoder (Epi-LSTM) framework. The latent representation resulting from the bottleneck of the autoencoder is then used for various downstream tasks.

## 2 Related work

Many methods have been proposed for genome annotation on the basis of epigenomics data sets [3], [4], [5], [6], [7],[8], [9], [10], [16]. As noted above, most existing methods produce celltype-specific annotations of activity in a given cell state. This includes a class joint annotation methods that aim to improve epigenome annotations by simultaneously annotating many cell types and sharing position-specific information between the annotations [17], [18], [19], [20], [21]. Such joint annotations can be more accurate, but still produce a separate annotation for each cell type.

The most related methods to ours aim to take data from all available cell types as input to produce a single pan-celltype (as opposed to celltype-specific) annotation, a task sometimes known as “stacked” annotation. Several methods have been proposed to produce discrete [22], [23] and continuous [13], [24] cell-type-agnostic representations.

## 3 Methods

### 3.1 Datasets

The epigenomic data pertaining to histone modifications and chromatin accessibility (ChIP-seq and DNase-seq) was downloaded from the Roadmap Epigenomics Consortium [1]. These experiments indicate the − log10 p-value of the enrichment at a particular genomic position compared to a reference track. Following Avocado and other frameworks before it like PREDICTD [24] and Segway [3], we use the arcsinh transformation (1) on the signal to stabilize the variance of these signals and lessen the effect of outliers.

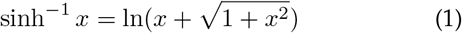

We consider four downstream classification tasks, namely: gene expression, promoter-enhancer interactions (PEIs), frequently interacting regions (FIREs), and replication timing.

We also conduct a downstream analysis pertaining to evolutionary conservation. Gene expression is characterized by RNA-Seq, a next-generation sequencing technology that measures the amount of RNA present in a biological sample. PEIs are revealed by ChIA-PET (Chromatin Interaction Analysis by Paired-End Tag Sequencing), a technique that quantifies genome-wide chromatin interactions. FIREs are located by Hi-C, a high-throughput sequencing technique to find fragments of nucleotide sequences. The replication of the genome is characterized by assays such as Repli-Seq [25], that separate the loci into early and late replicating segments. Evolutionary conservation at particular genomic sites is given by phyloP scores.

The data for the downstream tasks was obtained from resources pointed in and processed as mentioned in [13]. RNA-seq data for 57 cell types was obtained from [1]. We consider genes to be active if the logarithmic mean expression value across the gene is greater than 0.5 [26], [27]. These genes are assigned the label of 1 while the rest are assigned 0. The PEI’s used, are the ones that were used to train TargetFinder [28], which concur with ChIA-PET interactions. They were downloaded from the Repository [29] of TargetFinder, which includes the interaction data for four cell types namely, GM12878, HeLa-S3, IMR90 and K562. The data was preprocessed as given in [13]. FIRE score data was taken from the additional material of [30] for seven cell types, namely, TRO, H1, NPC, GM12878, MES, IMR90, and MSC. FIRE scores are given at 40kbp resolution and these are converted to binary indicators using a threshold of 0.5 [13]. Replication timing data was downloaded from [31] at a resolution of 40kbp. The phyloP scores are taken from the PHAST package [32] for multiple alignments of 99 vertebrate genomes to the human genome.

### 3.2 LSTM model

Our model employs an architecture which is based on the deep recurrent neural network (RNN) [33]. While regular neural networks expect fixed-size inputs and outputs, RNNs can handle inputs of arbitrary length, i.e. sequential, in nature. What distinguishes RNNs from other types of neural networks is that the output at any given step in the sequence is not just dependent on the input at that step but also on the entire history of inputs fed in the past. The RNN is able to achieve this dependency on preceding steps in the sequence by maintaining a hidden memory state that it carries over to future steps. As epigenomic data is sequential in nature, with short term interactions being highly prevalent, a sequential model like the RNN would be good choice for our embedding task.

RNNs rely on an extension of the Backpropagation algorithm called as the Backpropagation Through Time (BPTT), which links the derivatives between steps in the sequence. Often, a truncated version of BPTT is used that restricts the flow of gradients to specific steps in the past to reduce cost per parameter update. Although the Vanilla RNNs are quite powerful, they are subject to the problems of vanishing and exploding gradients [34], which makes it difficult for them to handle long-term dependencies. As an extension of the RNN architecture, Long Short-Term Memory (LSTM) is more effective at handling longer dependencies in steps, including the problems associated with gradient flow.

LSTMs were proposed as a solution to the vanishing gradient problem and consist of a cell which behaves as memory and three gates that act in tandem, namely, input gate, forget gate and output gate. The cell state of the LSTM keeps track of how much the preceding steps effect the steps to follow and the gates regulate this function. The input gate controls the amount a new input at a step effects the cell state, the forget gate regulates what portion of the cell state remains unaffected and the output gate discerns how much the cell state contributes to the output of the LSTM at a particular step. The equations that govern the working of the LSTM are given in (2).

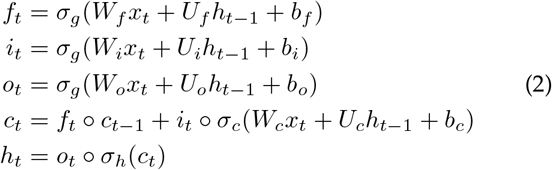

where the matrices *W_a_* and *h_a_* are the weights of the input and recurrent connections. The subscript _*a*_ can denote either the input (_*i*_), forget (_*f*_), output gate (_*o*_) or the cell state (_*c*_). While *h* denotes the hidden state, *o* denotes the output of the LSTM. *σ* refers to the sigmoid activation function and is the Hadamard product.

The LSTM maintains a representation of long term dependencies and, because of this ability, it serves to be a good candidate for modelling sequential data. Using these internal representations of the LSTM, we can recreate the original sequential input in a encoder-decoder framework [35], which forms the backbone of advances in fields like language modelling [36], speech recognition [37], sequence-to-sequence prediction [38] and neural machine translation (NMT) [39].

#### 3.2.1 Epi-LSTM Autoencoder Framework

The autoencoder forms the backbone of our proposed Epi-LSTM framework. The high dimensional sequential input assays are reconstructed via a low dimensional bottleneck using LSTMs coupled with a loss function that forces the output to be as close to the input as possible in euclidean space (Figure 2). The first stage of the framework is a LSTM that acts as an encoder. The encoder reads the input sequence and creates a fixed length low dimensional vector representation in the form of an embedding. The low dimensional representation at each position in the sequence is then treated as an annotation for that position. This annotation can then be used for various tasks related to the input sequence like recreating the input sequence as in auto-encoding or translation to a different sequence as in machine translation. The decoder uses the fixed length vector embedding as it’s initial hidden state and tries to recreate the original sequence. Along with the initial hidden state seed, the decoder, at each step, receives as input the output of the encoder and its cell state. It then uses these states to output the original input at each step. The Epi-LSTM is trained using the mean squared error (MSE) loss function (3) which facilitates reconstruction of the input. The errors are then back-propagated through the encoder-decoder framework and through steps in the sequence using BPTT.

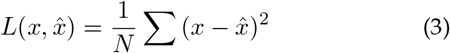

where *x* is the input vector at a particular step, 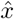 is the vector output of the decoder that step and *N* is the number of sequence steps in a a particular chunk of data.

**Fig. 2:**
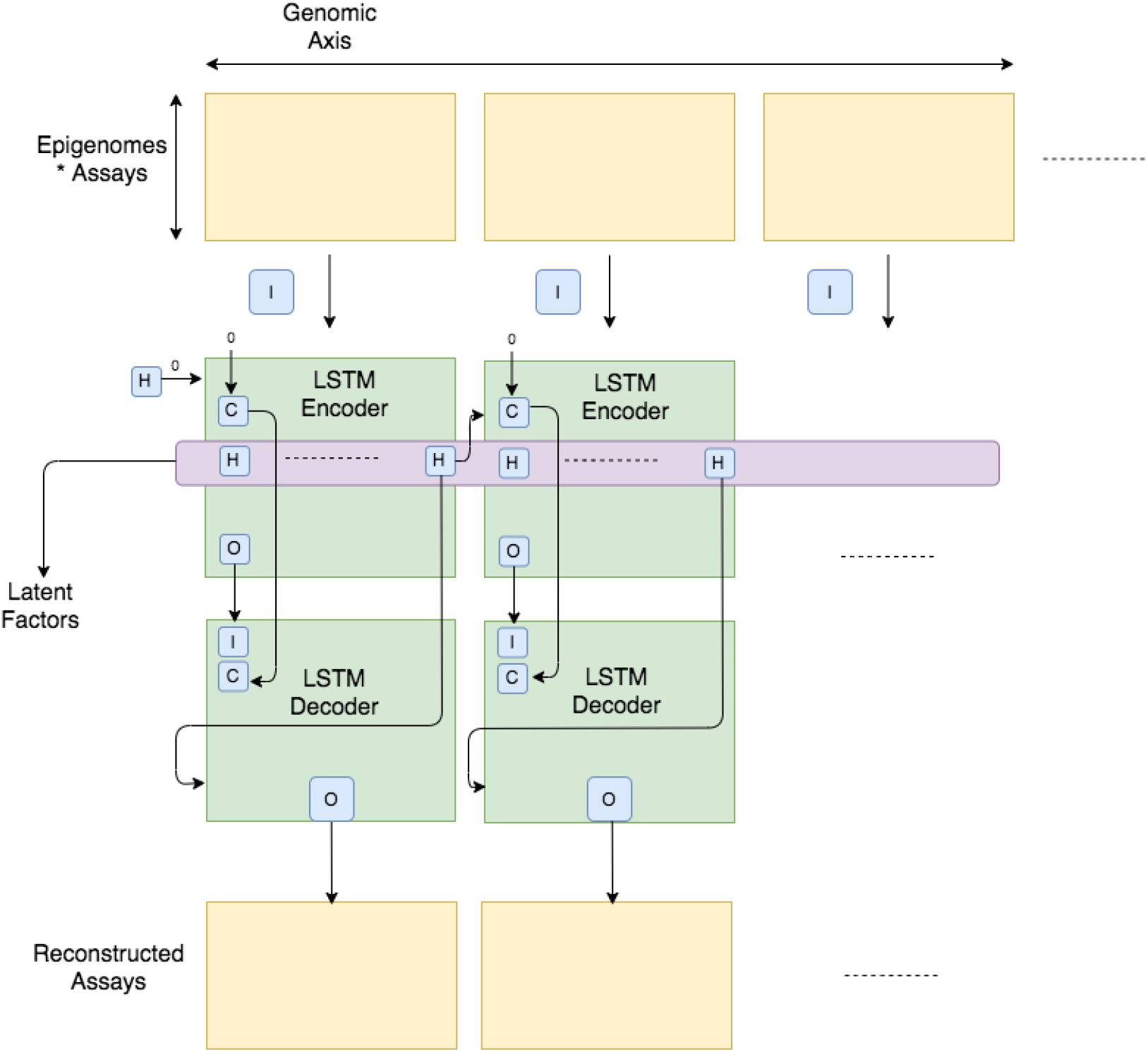
Epi-LSTM Framework. The Encoder and the Decoder are single-layer layer norm LSTMs. The assays that are serving as the input to the Encoder are arranged in a matrix of (Epigenomes*Assays) × Genome Length and fed in one Frame Length at a time. The Frame length is chosen as 100 to fit the model into memory.

The epigenomic data is fed into the Epi-LSTM in frames of 100 steps to fit the model into memory and to speed up training. As the epigenomic data that we used [1] had a resolution of 25, i.e., a data point for every 25 base pairs, Epi-LSTM is capable of dealing with 2500 positions in a given frame. At each step the epigenomic data vector can written as *EP_f_* = [*C*_1,*f*_ *A*_1,*f*_, …, *C*_1,*f*_ *A_K,f_*, …, *C*_2,*f*_ *A_K,f_*, …, *C_J,f_ A_K,f_*], with a total of *K* assays performed in *J* cell types. These epigenomic vectors are then transposed and stacked horizontally as *I_t_* = [(*EP*_1_)^*T*^, (*EP*_2_)*^T^*, …, (*EP_F_*)*^T^*], where *F* is frame length, *t* = 1, 2, …, *T* and *T* is the number of frames. For the Epi-LSTM, each column of *I_t_* is fed as input and the hidden and cell state are carried on to the next column (4).

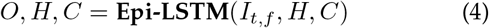

where *O* is the LSTM output, *H* is the hidden state and *C* is the cell state.

The hidden state of the encoder from the previous frame was carried over to the hidden state of the encoder in the next frame, whereas the cell state was reinitialized for each frame. This aids the model to not only carry information across frames but also maintain a fresh state particular to the current frame. The hidden state of the encoder at the end of a particular frame was fed into the decoder as its initial hidden state, whereas the cell state of the decoder at each step was the cell state of the encoder at that step and the input of the decoder at each step was the output of the encoder at that step (Figure 2). We arrived at these design choices by choosing the model that minimized reconstruction error following ablation experiments conducted with different variations of the model (supplementary material).

### 3.3 Comparison Methods

We compare the Epi-LSTM with Avocado’s Deep Tensor Factorization model [13], [40], REFINED+CNN and a majority baseline that always chooses the majority label in the data. Avocado is implemented as given in [13]. We applied REFINED [14] to the epigenomic data by treating the cell-type axis as the primary input to REFINED, i.e, we transposed the matrix *I_t_* as (*I_t_*)*^T^*, where the columns correspond to cell type-assay combinations which do not have explicit correlations and therefore can be treated as predictors in need of reordering, and rows are epigenomic data vectors. The transposed matrix would look like (*I_t_*)*^T^* = [*C*_1_*A*_1_, *C*_1_*A*_2_, …, *C*_1_*A_K_*, …, *C_J_ A_K_*], where each column corresponds to values of a cell type-assay combination for an entire frame length. This transposed matrix is fed as the input to REFINED, the output of which is reordered predictors that minimize the cost function using iterative hill climbing [14] (5). The cost function measures the absolute difference in the predictor distances among the new predictor locations to the estimated true distances [14].

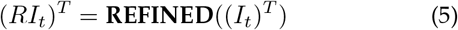

where (*RI_t_*)*^T^* has the columns reordered by REFINED. We further transpose this output to take it back to the original form *RI_t_*, with the columns as reordered epigenomic data vectors and feed it as the input to the CNN autoencoder.

### 3.4 Model Design Choices

We set the hidden size of our LSTMs to 110 (supplementary material.). The LSTM model was chosen to be unidirectional with one hidden layer in the presence of layer norm. By normalizing from all of the summed inputs to the neurons in a layer, layer normalization is shown to be very effective at stabilizing the hidden state dynamics in recurrent networks and can considerably reduce the training time [41]. It is therefore chosen as a default setting in our experiment. A variant, the Bidirectional LSTM, can be trained by using inputs not just up to the preset frame but by simultaneously feeding data forward and backward [42]. In addition to this, the idea of randomly dropping nodes and weights during training to prevent nodes from over adapting is called dropout [43]. The training phase forms an exponential number of “thinned” networks whose predictions are averaged during test time, which is known to help in regularization of the model and superior test performance. In order to mitigate hidden state saturation in the LSTM we used gradient clipping [44] and used the *softsign* activation [45] instead of alternatives such as *tanh* at all nodes. While Softsign has a similar range, it approaches its extremes much slower when compared to the hyperbolic tangent because it has quadratic rather than exponential tails. The hyperparameter and design choices were made after conducting ablation experiments which are elaborated in the supplementary material.

### 3.5 Training Procedure

The Epi-LSTM was built using the Python-based deep learning framework PyTorch [46] and trained on GeForce GTX 1080 Ti GPUs. The ADAM optimizer [47] was used for training due to its pervasive adoption in the machine learning community and the autoencoder was trained using the MSE loss function (3). Apart from the design choices mentioned in section 3.4, the other parameters were set to their default values in PyTorch while training.

### 3.6 Downstream Classification

The representation used to reconstruct the epigenomic data might be able to serve as a good basis for classification of genomic properties, as many of these properties manifest themselves in the epigenomic signals. Therefore, we test whether the position specific latent representation learned via unsupervised learning is useful for genomic tasks that the model was not trained to achieve such as classifying important genomic phenomena like gene expression, PEIs, replication timing and FIREs across cell types. For each task and cell type, we trained a XGBoost classifier [48] following [13], with a maximum of 5000 estimators and a maximum depth of 6 (supplementary material) on the representations obtained by training Epi-LSTM on the full set of ChIP-seq and DNase-seq assays in the Roadmap compendium [1]. XGBoost stands for eXtreme Gradient Boosting and is an implementation of gradient boosted decision trees designed for high performance with support for Stochastic and Regularized Gradient Boosting. In gradient boosting, new models that predict the residuals of previous models are combined to make the final prediction, using gradient descent to minimize the loss when combining models. We use n-fold cross-validation to perform and evaluate our training with *n* = 20 and an early stopping criterion to stop the training if the validation performance did not improve after 20 epochs. All other hyperparameters of XGBoost were kept at their default values. We used the metric of mean Average Precision (mAP), which is a standard metric in machine learning classification tasks, to evaluate our classification model. As the name suggests, mAP averages the precision values obtained at different levels of recall. The maximum precision value that can be achieved at a particular recall cutoff is calculated, and is repeated for all predefined recall levels. An average of these maximum values yields the mAP. Other classification metrics like Area under the Receiver Operating Characteristic (ROC)-curve (AuROC) and Accuracy are included in the supplementary material.

### 3.7 N-fold Cross-Validation

There are two main approaches for validating genomic classification models, cross-cell type and cross-chromosomal. Our classification model adopts the cross-chromosomal approach for validation by building a separate XGBoost classifier for each cell-type and partitioning the genome into *N* = 20 folds according to chromosomes (we pick 20 out of 22 chromosomes randomly). We do this because we are more interested in verifying the utility of the input Epi-LSTM representations than building one classifier that works across all cell-types. Note that this differs from a cross-cell type validation approach, which would be required if our aim was to use the model’s predictions on cell types without available data. Instead, we are interested in verifying that Epi-LSTM’s representations have the information required to recapitulate the known phenomena in the input cell types. Therefore, our validation approach is not prone to the overfitting pitfall mentioned in [49], which affects cross-cell type validation approaches.

## 4 Results

### 4.1 Representations capture many types of genomic activity

A good latent representation of the epigenome should contain all of the information about the regulatory state of each genomic locus. Therefore, to evaluate our representation, we asked whether known genomic phenomena are predictable from only the representation.

#### 4.1.1 Gene Expression

The expression of a gene as measured by techniques such as RNA-seq is influenced and therefore could be predicted by histone modifications as measured by ChIP-seq. Although it might be hard to interpret the exact relationship between the representations and the expression of a gene, it would be promising if they encode a complex genomic phenomena which is a result of varied interacting entities.

##### Classification

The mRNA abundance of a gene can either be expressed to several decimal places or only as binary, as to whether the gene is expressed or not. There have been many works that show the efficacy of using binary gene expression levels for classification [50], [51], [52], [53], [54], [55]. Following Avocado [13], [26], and [27], we set a threshold value of 0.5 on logarithmic scale of expressions. There are also other known ways of quantizing gene expression data like fitting a Gaussian mixture model to log expression levels as in [55] or discretization using sudden changes in sorted gene expressions as in [54].

Following [13], we predict gene expression by using representations from the promoter region. We find that the models built using the Epi-LSTM representations don’t perform as well as the Avocado representations in cell types where both perform very well, however, the Epi-LSTM representations perform better than Avocado in cell types where both perform poorly (Figure 3a). Both models perform significantly better than REFINED+CNN and the baseline (Figure 3a).

**Fig. 3:**
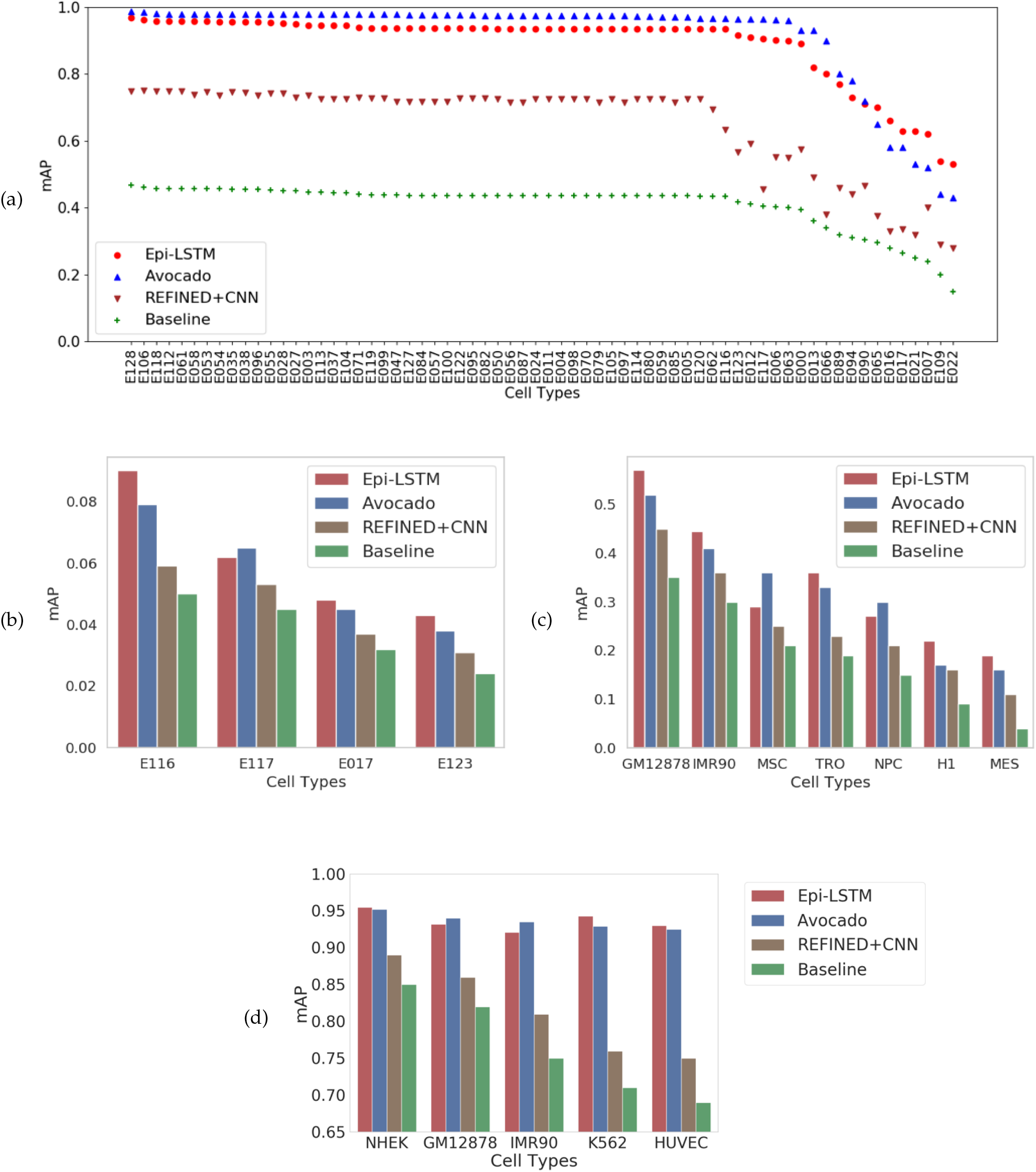
The average precision with which each method predicts gene expression, promoter-enhancer interactions, frequently interacting regions and replication timing as given by mAP is shown in Figures 3a, b, c and d respectively. The y-axis shows the mAP and the x-ticks refer to the cell types in all plots. The colour scheme is as follows: Epi-LSTM is shown in red, Avocado is shown in blue, REFINED+CNN is shown in brown and the Baseline is shown in green.

##### Hard-to-predict Cell Types

We found that Epi-LSTM achieves good performance even in hard-to-predict cell types. While Avocado outperforms the Epi-LSTM at predicting gene expression in many cell types, it does so in cell-types in which both the methods do extremely well in (> 0.95 mAP) and the difference in performance is marginal. In cell types that are harder to predict gene expression in (Aorta, Fetal Lung Fibroblasts, Blastocysts, Foreskin Fibroblasts, Neuronal Progenitor, Small Intestine), the Epi-LSTM outperforms Avocado (0.1 mAP higher on average). The performance of the majority baseline deteriorates considerably in these cell types as well, leading us to believe that it is easier to categorize gene expression in certain cell types compared to others. While Avocado does really well at predicting gene expression in many cell types, the Epi-LSTM performs more consistently across cell types, doing particularly well in the hard-to-predict cell types (Figure 3a).

#### 4.1.2 Promoter-Enhancer Interactions

Gene expression is also modulated by the promoter and enhancer elements interacting over relatively long distances. Promoter–enhancer interactions (PEIs) represent a subset of interactions of the chromatin which are key to the understanding of transcriptional regulation [56]. PEIs are vital for transcriptional regulation of an enhancer’s target gene, so much so that the target gene expression is observed to be affected by addition of PEI-disrupting insulators, lack of PEI-associated proteins, and gain of competing promoters [57]. Chromatin interactions are also known to be highly associated to gene co-expression rates [58]. Although these interactions can be captured by methods such as Hi-C [59], because these methods are expensive, it would be favourable to divine these from existing data that is much more easier to acquire. There have been methods that explore machine learning techniques for this purpose, like the one that uses representations derived from both the promoter and enhancer regions [28]. In our case we use representations derived from the window between the promoter and the enhancer elements and find that the models trained on the Epi-LSTM representations perform better than other methods on average across cell types, doing worse in some, while doing better in some others (Figure 3b). All methods have very low mAP, revealing the difficulty of this task. However, Epi-LSTM outperforms the other methods, indicating that Epi-LSTM representation contain the most information related to PE-interactions (Figure 3b).

#### 4.1.3 Chromatin Architecture

The three-dimensional folding of the chromatin orders the genome into compartments and creates spatial proximity between distant functional elements. The strength of interactions between pairs of loci in the genome can be obtained in a high throughput manner by techniques such as Hi-C [59], which is an extension of 3C, that is capable of locating genome-wide long range interactions. The genome is known to be partitioned into functional segments called topologically associated domains (TADs), where there is high interaction between the loci. TADs are a central feature of genome folding and were discovered in one of the first chromatin folding maps. It is believed that most of the mammalian genome is folded into globular domains of chromatin connected by boundaries which are linear [60]. Frequently interacting regions (FIREs) have been identified recently [30], which are regions generally present within TADs. FIREs are regions that live in between the anchor points of chromatin loops and are found to be tissue specific with higher enrichment near enhancers and super-enhancers. The task of predicting FIREs is posed as a classification task for each genomic locus and we find that models trained using the Epi-LSTM representations perform better on average across cell types compared to the other methods and the baseline (Figure 3c).

#### 4.1.4 Replication Timing

The genome replication within eukaryotic genomes is known to correlate with genome evolution, chromatin structure, and gene expression. It’s known that there exists a noticeable correlation between replication timing and the three-dimensional conformation of the chromatin and it is substantially stronger when compared to binding proteins and histone modifications [61]. The temporal and spatial characterization coupled with markedly altered epigenetic states of early-replicating when compared to late-replicating loci dictate the nature of chromosomal rearrangement in cells [62]. Although this suggests that replication timing might be controlled at the domain level, it is known to facilitate the organization of the epigenome as well [62] and therefore is a useful property to predict from a model that is trained on epigenomic data. The replication of the genome follows certain profiles as elaborated in [63], [64] which is useful for its characterization. We find that the model trained using the Epi-LSTM representations perform better than other methods in a majority of cell types (Figure 3d). Both Epi-LSTM and Avocado perform very well at this task, achieving *>*0.92 mAP on average (Figure 3d).

#### 4.1.5 Discussion

Epi-LSTM outperforms the other methods in a majority of cell-types on the tasks of PE-interactions, frequently interacting regions and replication timing. Particularly for PE-interactions, the mAP is quite low for all methods, showing the difficulty of predicting them from epigenomic representations. However, Epi-LSTM performs best of the methods, indicating that it includes the most information related to PE-interactions. While Avocado outperforms the Epi-LSTM at predicting gene expression in majority of cell types, the Epi-LSTM performs more consistently and does better in the hard-to-predict cell types.

### 4.2 Epi-LSTM captures evolutionarily conserved activity

To explore our representations further we performed visualizations related to salient information pertaining to genomic sites, that demonstrate that the model is indeed capturing interesting genomic properties. We find that our representation accurately identifies evolutionarily conserved regions in the genome. Genomic sequences of an evolutionary lineage can change due to random mutations and deletions in time over many generations. Sometimes chromosomal rearrangements may also promote them to recombine [65], [66]. Sequences in the genome that propagate despite such above mentioned forces, and therefore have lower mutation rates compared to the rest of the genome are said to be conserved [67]. Conservation manifests at coding and non-coding bases. Although highly conserved regions are known to have some functional importance, many highly conserved non-coding regions are poorly studied. Various parameters like genetic drift, population size, robustness of a sequence to mutation, and selection pressure affect the extent to which a sequence is conserved. PhyloP scores measure evolutionary conservation at particular genomic sites. Negative scores indicate evolutionary acceleration, which is faster than neutral drift and positive scores indicate evolutionary conservation, which is slower than neutral drift. These p-scores help us understand the nature of selection at chosen genomic positions. The absolute values of these scores stand for the −log p-values under a null hypothesis of neutral evolution. The 2D histogram of the phyloP scores with the feature values demonstrates that certain chosen representations are indicative of evolutionary conservation and acceleration (Figure 4).

**Fig. 4:**
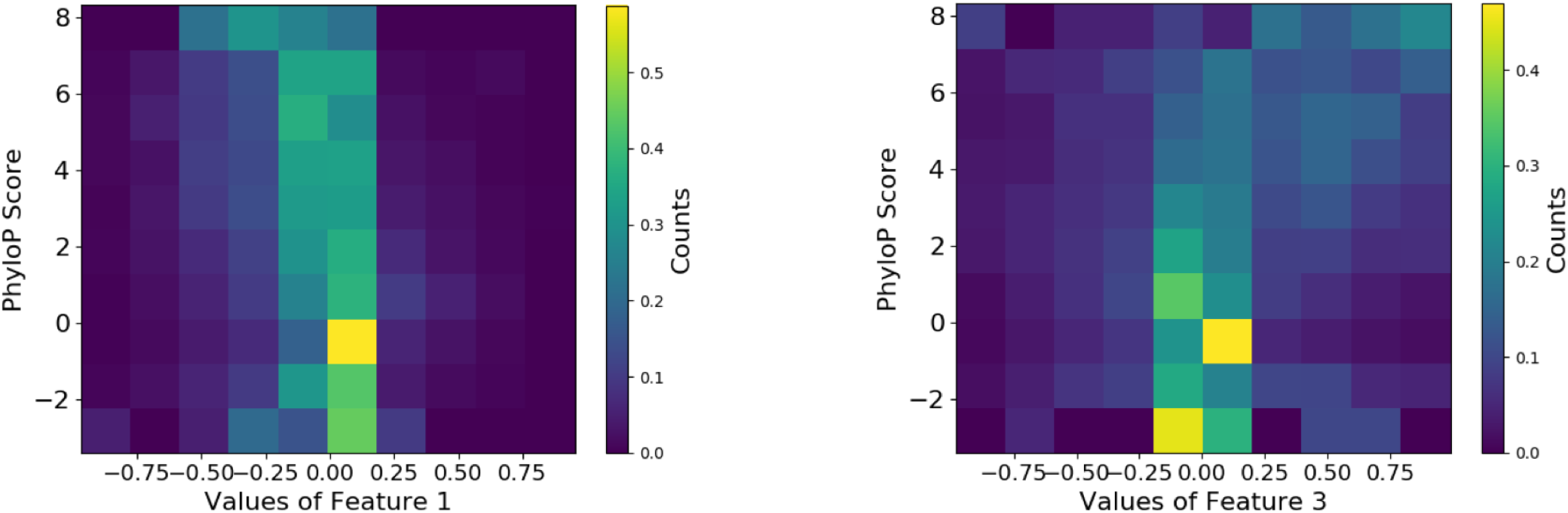
2D histogram of phyloP score and feature values. The x-axis bins the values of the chosen feature and the y-axis bins the values of the phyloP score. The color gradient denotes the strength of points in each bin after columnwise normalization.

### 4.3 Epi-LSTM representations correlate with phyloP score and GC content

To visualize the relationship between Epi-LSTM representations and conservation, we plot the correlation of the phyloP scores with our feature set and observe a positive correlation (Figure 5a). The GC-content refers to the percentage of nitrogenous bases in the DNA molecule that correspond to Guanine or Cytosine. There exists evidence that the GC content is higher for genes with longer coding regions i.e. the coding sequence length is proportional to GC content [68]. Genomic fragments with high GC content are also shown to have enhanced malleability, aiding nucleosome binding. Adding to this, compared to the noncoding regions of the genome, GC rich exons contain higher levels of DNA methylation [69]. Owing to the intimate relationship between coding regions, DNA methylation and GC content, we explore whether some sort of information about the GC content is captured in our representations. We plot the correlation between the GC-content and our feature vectors (Figure 5b) and observe a similar trend as with the p-score i.e. the GC-content shows an increasing correlation with our feature vectors (Figure 5b).

**Fig. 5:**
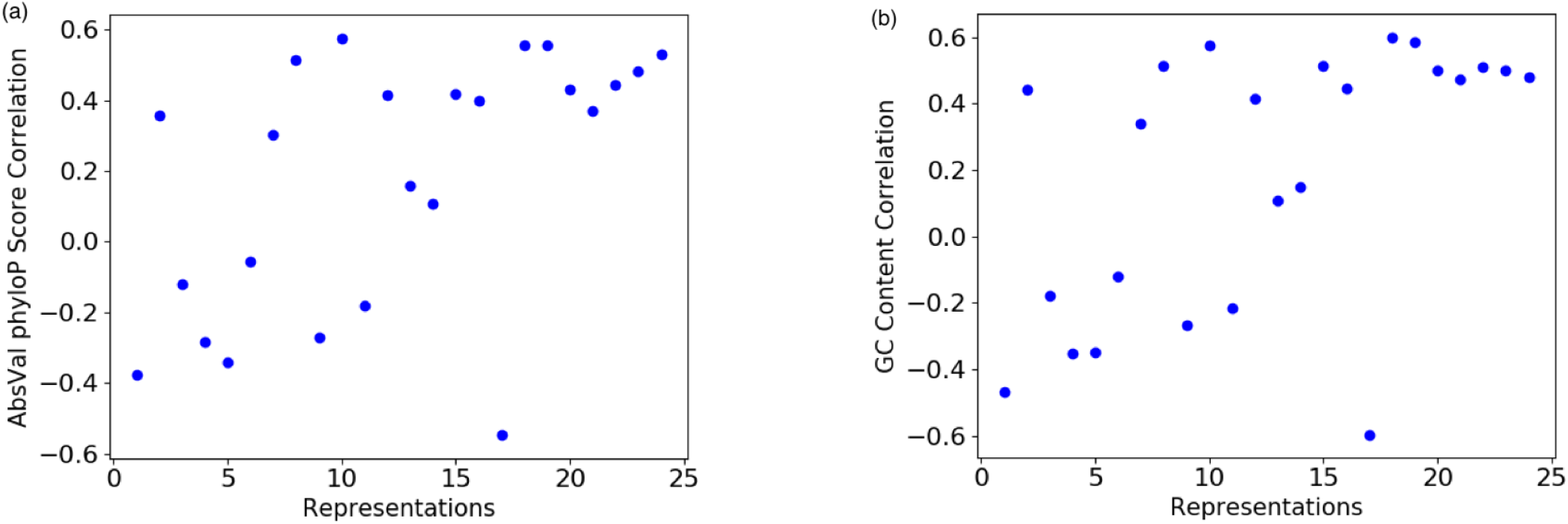
Correlation of Absolute value of phyloP score (a) and GC content (b) with Epi-LSTM representations. The x-axis shows the representations and the y-axis displays the correlation. The blue dots represent the correlations corresponding to the given representation.

### 4.3 wEpi-LSTM provides a smooth representation along the genomic axis

Smoothness of the representations across positions in the genome gives us an indication of how well the model is able to take into account the similarity of nearby positions and how the representations change as one moves further away along the genomic axis. Smooth representations match our biological expectation that contiguous genomic elements have a common function. In contrast, rapid fluctuations in representation space indicate biologically implausible fluctuations in activity. This knowledge can then be used to interpret any observed departure from smoothness either due to functional variability or experimental artifacts. Setting a standard for how smooth the representations need to be on average across the genome allows us to not only ensure closeness of nearby positions in representation space but also characterize genomic properties that depart from this standard. Similar to the standard definition of smoothness that is characterized by the continuity of the surface made up of the points in a higher dimension, we measure the smoothness of the points in euclidean space by calculating a surrogate R-squared value which gives the proportion of the variance of the representations at position *x* + *k* explained by the representations at position *x*, averaged across the genome for different values of *x* and tabulated for varying values of *k*. Owing to the sequential nature of Epi-LSTM, we expect the representations obtained to be smoother across the genome when compared to other methods and we do in fact observe the same. To demonstrate this, we plot the aforementioned R-squared metric between pairs of positions for different distances and average it across the genome (Figure 6) and it can be seen that the Epi-LSTM representations are smoother than representations from Avocado and REFINED+CNN over a large portion of the genome.

**Fig. 6:**
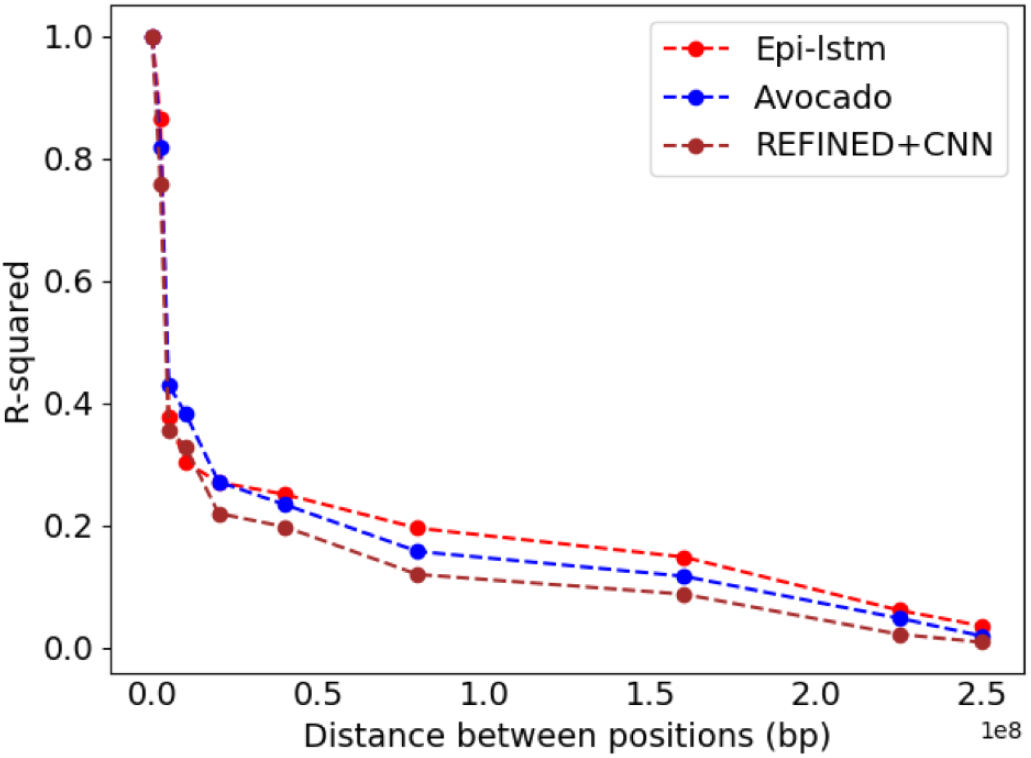
Regression performance of epigenomic reconstruction using the representations of different methods as given by R-squared. The x-axis represents the difference between the positions in the genome in base pairs. The y-axis shows the R-squared for pairs of positions for varied distances averaged across the genome. The legend shows the different methods. The colour scheme is as follows: The Epi-LSTM is coloured red, Avocado is coloured blue, and REFINED+CNN is coloured brown.

## 5 Conclusion

In this work we have proposed a deep LSTM autoencoder called Epi-LSTM that uses epigenomic data from all cell types to form latent representations that can be used as pan-celltype position specific annotations. We demonstrate that these annotations are able to classify a variety of genomic phenomena while at the same time capturing information about important genome wide properties such as evolutionary conservation and GC content. Exploring relative feature importance and other powerful sequential models like Transformers [70] are interesting avenues for extension and remain as part of future work.

The primary contribution of this work is the application of a recurrent LSTM neural network structure to this problem. This sequential structure is a natural fit to genomics data sets because of the sequential nature of the genome. For that reason, sequential models such as hidden Markov models are widely used in genomics. We found that, likely due to the sequential nature of the model, its latent representation is smoother along the genomic dimension that a model relying purely on embeddings. The code repository for this project, including training, evaluation and data handling can be found at [71].

This work produces a pan-celltype representation to the epigenome, in contrast to related SAGA methods. SAGA methods are cell type-specific; they produce a separate annotation for each cell type. A pan-celltype representation is greatly needed because it makes it easy to understand the function and evolution of a genomic locus. Given that most other widely-used genome annotations (such of genes) are pan-celltype, we expect that pan-celltype epigenome annotations will soon become widely used in genomic analysis.

## Appendix A Supplementary Meterial

The supplementary material (attached as a separate file) contains: additional experiments, ablation results, hyper-paramter search plots, additional classification metrics and training and testing time plots.

---

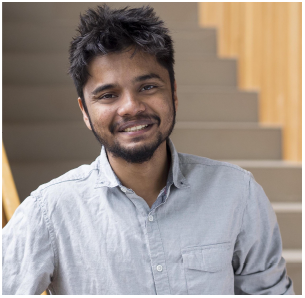
**Kevin B. Dsouza** completed his BTech in Electronics and Communication Engineering from National Institute of Technology Karnataka, India followed by his MASc in Electrical and Computer Engineering from The university of British Columbia (UBC) where he worked on machine learning techniques for next generation wireless systems. Currently, he is a PhD student at UBC where he works on computational genomics jointly under the information theory group at UBC and computational biology group at The Simon Fraser University. His work focuses on building machine learning tools to aid in the understanding of structural and functional genomic data.

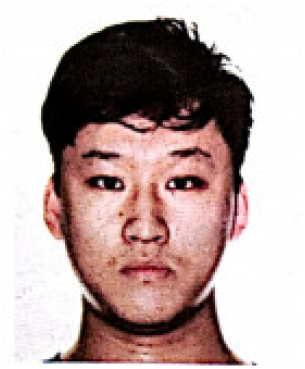
**Adam Y. Li** is currently a Master’s student in the Bioinformatics and Systems Biology program at the University of Amsterdam and Vrije Universiteit Amsterdam. He received his Honors Bachelor degree in Physics & Mathematics from the University of Toronto. His focuses are on the applications of computer and data science in the fields of bioinformatics and aging.

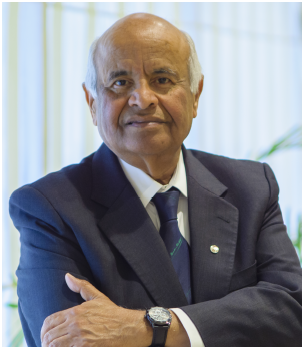
**Vijay K. Bhargava** (S’70, M’74, SM’82, F’92, LF’13) was born in Beawar, India. He obtained BASc, MASc and PhD degrees from Queen’s University at Kingston. He is a Professor in the Department of Electrical and Computer Engineering at the University of British Columbia in Vancouver. Vijay is a Fellow of the IEEE, The Royal Society of Canada, The Canadian Academy of Engineering and the Engineering Institute of Canada. Vijay has served as Director of Region 7 (1992, 1993), President of the Information Theory Society (2000), President of the IEEE Communications Society (2012, 2013) and as the Director for Division III (2018). He has served as an Associate Editor of the IEEE Transactions on Communications and as the Editor-in-Chief of the IEEE Transactions on Wireless Communications.

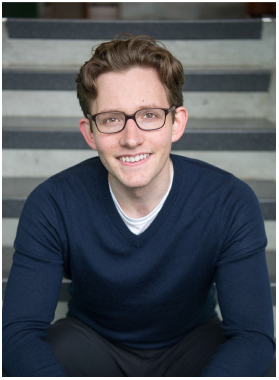
**Maxwell W. Libbrecht** is an Assistant Professor in Computing Science at Simon Fraser University. He received his PhD in 2016 from the Computer Science and Engineering department at University of Washington, advised by William Noble and Jeff Bilmes. He received his undergraduate degree in Computer Science from Stanford University, where he did research with Serafim Batzoglou. His research focuses on developing machine learning methods applied to high-throughput genomics data sets.

## Notes

### Competing Interest Statement

The authors have declared no competing interest.

## References

[1] The Roadmap Epigenomics Mapping Consortium. [Online]. Available: http://www.roadmapepigenomics.org/

[2] Encyclopedia of DNA Elements. [Online]. Available: https://www.encodeproject.org/

[3] M. Hoffman, O. J. Buske, J. Wang, Z. Weng, J. A. Bilmes, & W. S. Noble. Unsupervised pattern discovery in human chromatin structure through genomic segmentation. Nature methods, 9(5), 473–476. 2012.

[4] J. Ernst, & M. Kellis. ChromHMM: automating chromatin-state discovery and characterization. Nature methods, 9(3), 215. 2012.

[5] N. Day, A. Hemmaplardh, R. E. Thurman, J. A. Stamatoyannopoulos, & W. S. Noble. Unsupervised segmentation of continuous genomic data. Bioinformatics, 23(11), 1424–1426. 2007.

[6] J. L. Larson, C. Huttenhower, J. Quackenbush, & GC. Yuan. A tiered hidden Markov model characterizes multi-scale chromatin states. Genomics 102(1), 1–7. 2013.

[7] KA. Sohn, J. WK. Ho, D. Djordjevic, HH. Jeong, P. J. Park, & J. H. Kim. hiHMM: Bayesian non-parametric joint inference of chromatin state maps. Bioinformatics 31(13), 2066–2074. 2015.

[8] A. Mammana, & HR. Chung. Chromatin segmentation based on a probabilistic model for read counts explains a large portion of the epigenome. Genome biology 16(1), 151. 2015.

[9] J. Zhou, & O. G. Troyanskaya. Probabilistic modelling of chromatin code landscape reveals functional diversity of enhancer-like chromatin states. Nature communications, 7, 10528. 2016.

[10] S. G. Coetzee, Z. Ramjan, H. Q. Dinh, B. P. Berman, & D. J. Hazelett. Statehub-statepaintr: rapid and reproducible chromatin state evaluation for custom genome annotation. BioRxiv, 127720. 2017.

[11] S. Hochreiter, & J. Schmidhuber. Long short-term memory. Neural computation, 9(8), 1735–1780. 1997.

[12] H. I. H. Chen, Y. C. Chiu, T. Zhang, S. Zhang, Y. Huang, & Y. Chen. GSAE: an autoencoder with embedded gene-set nodes for genomics functional characterization. BMC systems biology, 12(8), 45–57. 2018.

[13] J. Schreiber, T. Durham, J. Bilmes, & W. S. Noble. Multi-scale deep tensor factorization learns a latent representation of the human epigenome. BioRxiv, 364976. 2019.

[14] O. Bazgir, R. Zhang, S. R. Dhruba, R. Rahman, S. Ghosh, & R. Pal. Representation of features as images with neighborhood dependencies for compatibility with convolutional neural networks. Nature communications, 11(1), 1–13. 2020.

[15] X. J. Mao, C. Shen, & Y. B. Yang. Image restoration using convolutional auto-encoders with symmetric skip connections. arXiv preprint arXiv:1606.08921. 2016.

[16] B. Chen, N. S. Kenari, & M. W. Libbrecht. Continuous chromatin state feature annotation of the human epigenome. bioRxiv, 473017. 2018.

[17] Y. Zhang, L. An, F. Yue, & R. C. Hardison. Jointly characterizing epigenetic dynamics across multiple human cell types. Nucleic acids research 44(14), 6721–6731. 2016.

[18] Y. Zhang, & R. C. Hardison. Accurate and reproducible functional maps in 127 human cell types via 2D genome segmentation. Nucleic acids research 45(17), 9823–9836. 2017.

[19] J. Biesinger, Y. Wang, & X. Xie. Discovering and mapping chromatin states using a tree hidden Markov model. In BMC bioinformatics, 14(S5), S4, BioMed Central. 2013.

[20] M. W. Libbrecht, M. M. Hoffman, J. A. Bilmes, & W. S. Noble. Entropic graph-based posterior regularization: Extended version. In Proceedings of the International Conference on Machine Learning. 2015.

[21] M. W. Libbrecht, F. Ay, M. M. Hoffman, D. M. Gilbert, J. A. Bilmes, & W. S. Noble. Joint annotation of chromatin state and chromatin conformation reveals relationships among domain types and identifies domains of cell-type-specific expression. Genome research, 25(4), 544–557. 2015.

[22] M. M. Hoffman, J. Ernst, S. P. Wilder, A. Kundaje, R. S. Harris, M. Libbrecht, … & R. C. Hardison. Integrative annotation of chromatin elements from ENCODE data. Nucleic acids research, 41(2), 827–841. 2013.

[23] M. W. Libbrecht, O. L. Rodriguez, Z. Weng, J. A. Bilmes, M. M. Hoffman, & W. S. Noble. A unified encyclopedia of human functional DNA elements through fully automated annotation of 164 human cell types. Genome biology, 20(1), 180. 2019.

[24] T. J. Durham, M. W. Libbrecht, J. J. Howbert, J. Bilmes, & W. S. Noble. PREDICTD parallel epigenomics data imputation with cloud-based tensor decomposition. Nature communications, 9(1), 1402. 2018.

[25] C. Marchal, T. Sasaki, D. Vera, K. Wilson, J. Sima, J. C. Rivera-Mulia, … & D. M. Gilbert. Genome-wide analysis of replication timing by next-generation sequencing with E/L Repli-seq. Nature protocols, 13(5), 819. 2018.

[26] N. Friedman, M. Linial, I. Nachman, & D. Pe’er. Using Bayesian networks to analyze expression data. Journal of Computer Biology, 7(3–4), 601–620. 2000.

[27] D. Pe’er, A. Regev, G. Elidan, & N. Friedman. Inferring subnetworks from perturbed expression profiles. Bioinformatics, 17(1), S215–224. 2001.

[28] S. Whalen, R. M. Truty, & K. S. Pollard. Enhancer–promoter interactions are encoded by complex genomic signatures on looping chromatin. Nature genetics, 48(5), 488. 2016.

[29] The TargetFinder Repository. TargetFinder. [Online]. Available: https://github.com/shwhalen/targetfinder

[30] A. D. Schmitt, M. Hu, I. Jung, Z. Xu, Y. Qiu, C. L. Tan, … & B. Ren. A compendium of chromatin contact maps reveals spatially active regions in the human genome. Cell reports, 17(8), 2042–2059. 2016.

[31] Replication Timing data. ReplicationDomain. [Online]. Available: http://www.replicationdomain.or

[32] The PHAST Package. PHAST. [Online]. Available: http://compgen.bscb.cornell.edu/phast/

[33] J. L. Elman. Finding structure in time. Cognitive Science, 14:179–211. 1990.

[34] R. Pascanu, T. Mikolov, & Y. Bengio. On the difficulty of training recurrent neural networks. In International conference on machine learning, 1310–1318. 2013.

[35] K. Cho, B. Van Merriënboer, C. Gulcehre, D. Bahdanau, F. Bougares, H. Schwenk, & Y. Bengio. Learning phrase representations using RNN encoder-decoder for statistical machine translation. arXiv preprint arXiv:1406.1078. 2014.

[36] I. Sutskever, O. Vinyals, & Q. V. Le. Sequence to sequence learning with neural networks. In Advances in neural information processing systems, 3104–3112. 2014.

[37] L. Lu, X. Zhang, K. Cho, & S. Renals. A study of the recurrent neural network encoder-decoder for large vocabulary speech recognition. In Sixteenth Annual Conference of the International Speech Communication Association. 2015.

[38] T. Young, D. Hazarika, S. Poria, & E. Cambria. Recent trends in deep learning based natural language processing. ieee Computational intelligenCe magazine, 13(3), 55–75. 2018.

[39] K. Cho, B. Van Merriënboer, D. Bahdanau, & Y. Bengio. On the properties of neural machine translation: Encoder-decoder approaches. arXiv preprint arXiv:1409.1259. 2014.

[40] The Avocado project, The Multi-scale deep tensor factorization Model. [Online]. Available: https://noble.gs.washington.edu/proj/avocado/

[41] J. Lei Ba, J. R. Kiros, & G. E. Hinton. Layer normalization. arXiv preprint, arXiv:1607.06450. 2016

[42] M. Schuster, & K. K. Paliwal. Bidirectional recurrent neural networks. IEEE transactions on Signal Processing, 45(11), 2673–2681. 1997.

[43] N. Srivastava, G. Hinton, A. Krizhevsky, I. Sutskever, & R. Salakhutdinov. Dropout: a simple way to prevent neural networks from overfitting. The journal of machine learning research, 15(1), 1929–1958. 2014.

[44] R. Pascanu, T. Mikolov, & Y. Bengio. On the difficulty of training recurrent neural networks. In International conference on machine learning, 1310–1318. 2013.

[45] X. Glorot, & Y. Bengio. Understanding the difficulty of training deep feedforward neural networks. In Proceedings of the thirteenth international conference on artificial intelligence and statistics, 249–256. 2010.

[46] PyTorch. [Online]. Available: https://pytorch.org/

[47] D. P. Kingma, & J. Ba. Adam: A method for stochastic optimization. arXiv preprint arXiv:1412.6980. 2014.

[48] T. Chen, & C. Guestrin. Xgboost: A scalable tree boosting system. In Proceedings of the 22nd acm sigkdd international conference on knowledge discovery and data mining, 785–794. 2016.

[49] J. Schreiber, R. Singh, J. Bilmes, & W. S. Noble. A pitfall for machine learning methods aiming to predict across cell types. Genome biology, 21(1), 1–6. 2020.

[50] S. Tuna, & M. Niranjan. Classification with binary gene expressions. Journal of Biomedical Science and Engineering, 2(6), 390–399. 2009.

[51] S. Draghici, P. Khatri, A. C. Eklund, & Z. Szallasi. Reliability and reproducibility issues in DNA mi-croarray measurements. Trends in Genetics, 22(2), 101–109. 2006.

[52] D. Geman, C. d’Avignon, D. Q. Naiman, & R. L. Winslow. Classifying gene expression profiles from pairwise mRNA comparisons. Statistical Applications in Genetics and Molecular Biology, 3. 2004.

[53] S. Tuna, & M. Niranjan. Inference from low precision transcriptome data representation. Journal of Signal Processing Systems, 58(3), 267–279. 2009

[54] I. Shmulevich, & W. Zhang. Binary analysis and optimization-based normalization of gene expression data. Bioinformatics, 18(4), 555–565. 2002.

[55] X. Zhou, X. Wang, & E. R. Dougherty. Binarization of microarray data on the basis of a mixturemodel. Molecular Cancer Therapeutics, 2(7), 679–684. 2003.

[56] A. Mora, G. K. Sandve, O. S. Gabrielsen, & R. Eskeland. In the loop: promoter–enhancer interactions and bioinformatics. Briefings in bioinformatics, 17(6), 980–995. 2016.

[57] I. Krivega, & A. Dean. Enhancer and promoter interactions—long distance calls. Current opinion in genetics & development, 22(2), 79–85. 2012.

[58] X. Dong, C. Li, Y. Chen, G. Ding, & Y. Li. Human transcriptional interactome of chromatin contribute to gene co–expression. BMC genomics, 11(1), 704. 2010.

[59] N. L. Van Berkum, E. Lieberman-Aiden, L. Williams, M. Imakaev, A. Gnirke, L. A. Mirny, … & E. S. Lander. Hi-C: a method to study the three-dimensional architecture of genomes. JoVE (Journal of Visualized Experiments), (39), e1869. 2010.

[60] J. A. Beagan, & J. E. Phillips–Cremins. On the existence and functionality of topologically associating domains. Nature Genetics, 1–9. 2020.

[61] N. Rhind, & D. M. Gilbert. DNA replication timing. Cold Spring Harbor perspectives in biology, 5(8), a010132. 2013.

[62] Q. Du, S. A. Bert, N. J. Armstrong, C. E. Caldon, J. Z. Song, S. S. Nair, … & W. Qu. Replication timing and epigenome remodelling are associated with the nature of chromosomal rearrangements in cancer. Nature communications, 10(1), 1–15. 2019.

[63] T. Ryba, I. Hiratani, J. Lu, M. Itoh, M. Kulik, J. Zhang, … & D. M. Gilbert. Evolutionarily conserved replication timing profiles predict long-range chromatin interactions and distinguish closely related cell types. Genome research, 20(6), 761–770. 2010.

[64] V. Dileep, F. Ay, J. Sima, D. L. Vera, W. S. Noble, & D. M. Gilbert. Topologically associating domains and their long-range contacts are established during early G1 coincident with the establishment of the replication-timing program. Genome research, 25(8), 1104–1113. 2015.

[65] M. Kimura. Evolutionary rate at the molecular level. Nature, 217(5129), 624–626. 1968.

[66] J. L. King, & T. H. Jukes. Non-darwinian evolution. Science, 164(3881), 788–798. 1969.

[67] M. Kimura, & T. Ohta. On some principles governing molecular evolution. Proceedings of the National Academy of Sciences, 71(7), 2848–2852. 1974.

[68] U. Pozzoli, G. Menozzi, M. Fumagalli, M. Cereda, G. P. Comi, R. Cagliani, … & M. Sironi. Both selective and neutral processes drive GC content evolution in the human genome. BMC evolutionary biology, 8(1), 99. 2008.

[69] S. Gelfman, & G. Ast. When epigenetics meets alternative splicing: the roles of DNA methylation and GC architecture. Epigenomics, 5(4), 351–353. 2013.

[70] A. Vaswani, N. Shazeer, N. Parmar, J. Uszkoreit, L. Jones, A. N. Gomez, … & I. Polosukhin. Attention is all you need. In Advances in neural information processing systems, 5998–6008. 2017.

[71] LSTM Model for the Epigenome. [Online]. Available: https://github.com/kevinbdsouza/latentGenome

