## Supplementary Material for "Latent representation of the human pan-celltype epigenome through a deep recurrent neural network"

### Supplementary Material: IEEE Paper-TCBB-2020-05-0292

This supplementary material contains additional experiments, ablation results, hyperparamter search plots, additional classification metrics and training and testing time plots.

#### I. Gene Expression Regression

We wanted to check if performing regression on the gene expression values gives us some more insight into the utility of the representations. We use RNA-seq expression values from [1] and perform regression on the logarithmic expression values using a neural network. The average R-squared values obtained for the different cell types shows a very similar trend as the classification result (Figure 1). Epi-LSTM performs similarly to Avocado in a majority of cell types and better than Avocado in certain hard-to-predict cell types. Both methods perform much better than REFINED+CNN and the baseline.

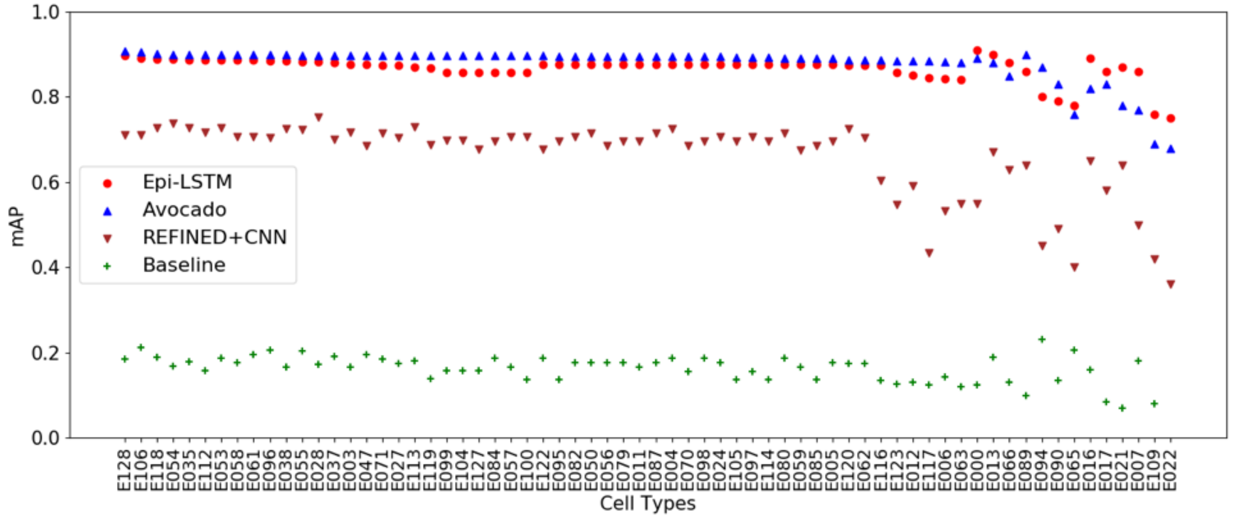


Fig. 1: Regression performance of predicting gene expression as given by R-squared. The y-axis shows the R-squared and the x-ticks refer to the cell types. The colour scheme is as follows: Epi-LSTM and Avocado are represented by red circles and blue triangles respectively, REFINED+CNN and the Baseline are represented by brown inverted triangles and green plusses respectively.

#### II. Ablation Results

The Epi-LSTM uses an autoencoder backbone to form representations. These representations are then used as an input the XGBoost classifier to predict important genomic phenomena. In order to check whether the LSTM and XGBoost are reasonable choices for their given tasks, we conduct two experiments, namely: (a) autoencoder ablation and (b) classifier ablation.

##### *A. Autoencoder Ablation*

In order to check how different type of autoencoder architectures perform at the task of reconstructing the epigenomic input, we conduct ablation experiments by replacing the LSTM autoencoder with three autoencoders: (a) Recurrent Neural Network (RNN) autoencoder, (b) Convolutional Neural Network (CNN) autoencoder, and (c) fully connected autoencoder (Figure 2b). For the RNN and the fully connected autoencoders we set number of hidden layers as 1 and the hidden size as 110, the same as the Epi-LSTM.

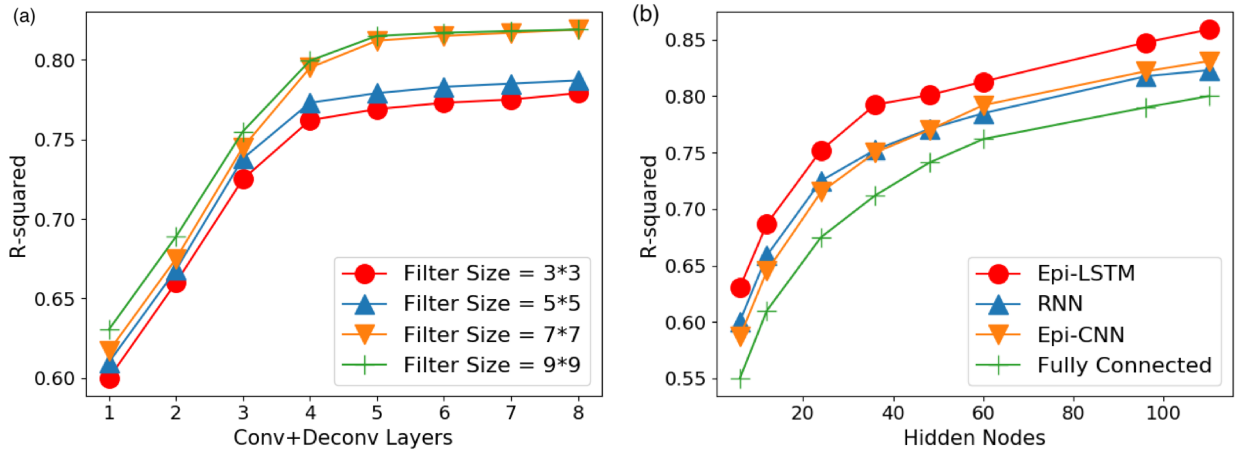


Fig. 2: (a) Regression performance of epigenomic reconstruction using CNN autoencoder as measured by R-squared. The y-axis shows the R-squared and the x-axis shows the number of Conv and Deconv layers used. The legend refers to the filter size. The number of hidden nodes at the innermost Conv layer is set to 110. (b) Regression performance of epigenomic reconstruction with different autoencoders as measured by R-squared. The y-axis shows the R-squared and the x-axis shows the number of hidden nodes used. The colour scheme is as follows: The Epi-LSTM is coloured red, RNN is coloured blue, Epi-CNN is coloured saffron and the fully connected is coloured green.

For the CNN autoencoder, we conduct a further search for the optimal values of filter size and number of convolutional and deconvolutional layers. We follow the CNN autoencoder architecture of [2] and use the same number of convolutional and deconvolutional layers. We search for the optimal filter size and Conv+Deconv layers (Figure 2a), and find the elbow at 5 layers and therefore set the number of Conv and Deconv layers as 5 each. We couple these with filters of filter size $7 * 7$, as they perform close to best with reduced expressive power. We set the number of hidden nodes at the innermost Conv layer as 110 and call this CNN autoencoder “Epi-CNN”.

R-squared for epigenomic reconstruction plotted against the number of hidden nodes for different autoencoders (Figure 2b) reveals that the Epi-LSTM performs best followed by the RNN and the Epi-CNN. The fully connected autoencoder performs the worst.

##### *B. Classifier Ablation*

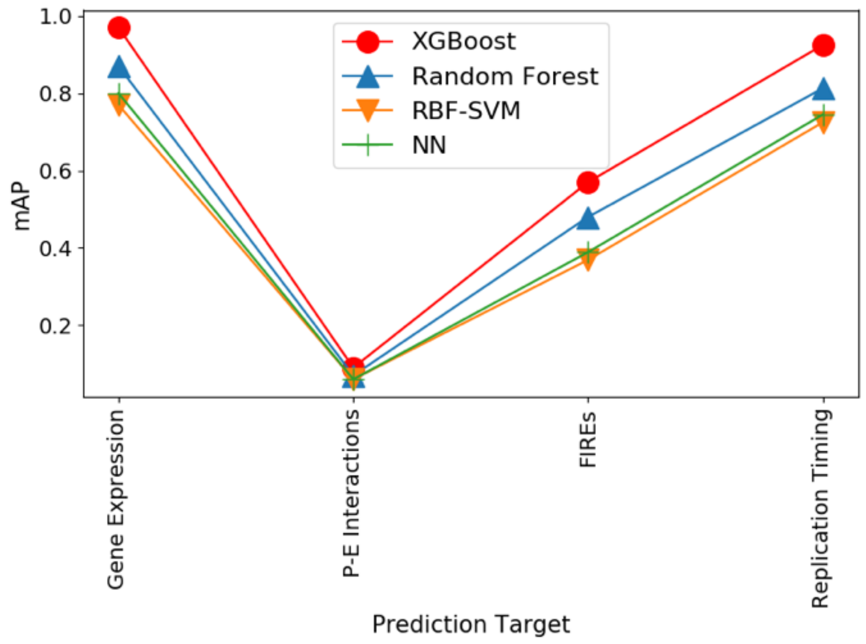


Fig. 3: The average precision for downstream genomic tasks using different classifiers as measured by mAP. The y-axis shows the mAP and the x-ticks show the downstream genomic tasks. The legend refers to the classifiers. The colour scheme is as follows: XGBoost is coloured red, Vanilla Random Forest is coloured blue, RBF-SVM is coloured saffron and the NN is coloured green.

In order to check whether XGBoost is the optimal choice for the task of downstream classification, we conduct an ablation study by replacing XGBoost with different classifiers, namely: (a) vanilla Random Forest, (b) Radial Basis Function Kernel Support Vector Machine (RBF-SVM), and (c) Neural Network (NN) classifier.

We set the parameters of the vanilla random forest the same as the gradient boosted XGBoost random forest. The RBF-SVM has two main parameters, namely: (a) $C$: This acts like a regularization parameter in the RBF-SVM. It provides a trade off between maximizing the decision function margin and training accuracy. A lower value of $C$ encourages a bigger margin, at the cost of correct classification. A bigger value of C accepts a smaller margin, if the decision function is good at classification. (b) $\gamma$: This corresponds to the inverse of the radius of influence of training samples. We find the optimal values for these two parameters using grid search and set them as 1 and 0.1 respectively. For the NN classifier, we set the number of hidden layers as 1 and number of hidden nodes as 32 after performing a grid search.

mAP plotted for the different genomic tasks using different classifiers (Figure 3) reveals that XGBoost is indeed a better classifier than the vanilla random forest and random forest in general is a better classifier than the RBF-SVM and NN.

#### III. Hyperparameter Search

We conduct hyperparameter search for the Epi-LSTM as well as the XGBoost classifier. The ablations for the Epi-LSTM include: (a) model architecture variations, (b) model functional variations, (c) number of hidden layers and nodes, and (d) learning rate. The main parameters for XGBoost are: (a) maximum depth and (b) maximum number of estimators.

##### *A. Epi-LSTM*

We define four variations for the Epi-LSTM architecture.

- Variation 1: Seed the last encoder hidden state of one frame length as the input for the encoder hidden state of the next frame length. Reinitialize the encoder cell state.
- Variation 2: Reinitialize both the encoder hidden and cell state.
- Variation 3: Seed the last encoder hidden state as the input for the decoder hidden state. At each step, the decoder cell state is the encoder cell state.
- Variation 4: At each step, both the decoder hidden and cell state are the encoder hidden and cell state respectively.

R-squared for epigenomic reconstruction plotted against the number of hidden nodes for different combinations of these variations (Figure 4) shows that Variation1+3, corresponding to the Epi-LSTM, performs the best among all combinations.

The Epi-LSTM uses single layer, unidirectional layer norm LSTMs without dropout. To check the effect of using dropout, removing layer norm, adding an additional layer or direction, we plotted the

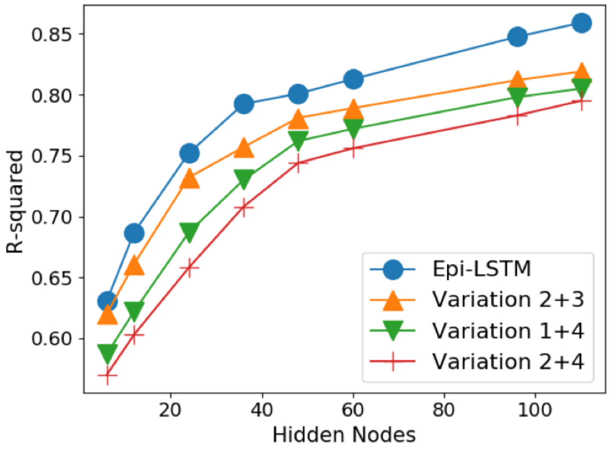


Fig. 4: Regression performance of epigenomic reconstruction for different number of hidden nodes with combinations of model variations as measured by R-squared. The y-axis shows the R-squared and the x-axis shows the number of hidden nodes. The legend refers to the combination of variations.

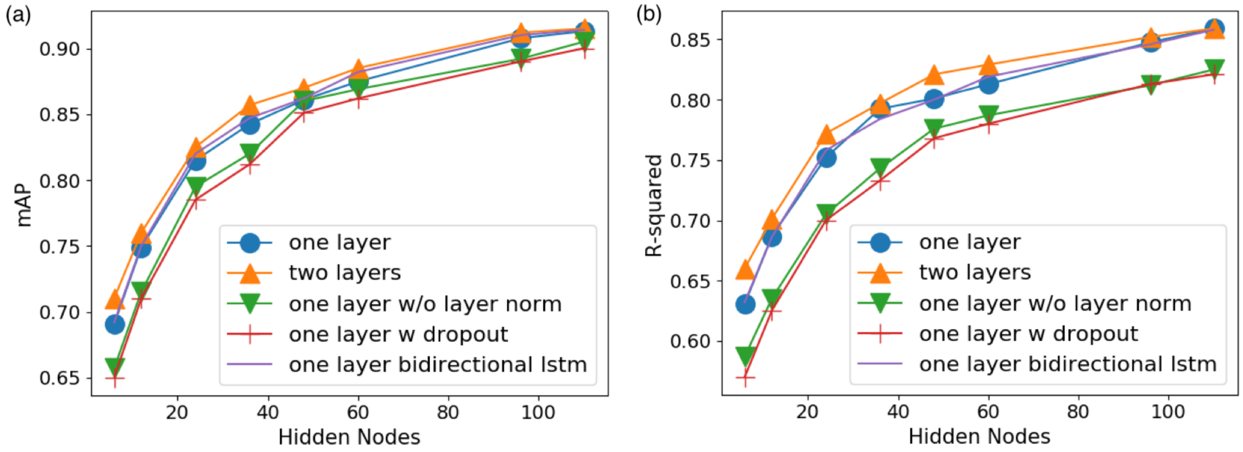


Fig. 5: The average precision for predicting gene expression as measured by mAP (a) and regression performance of epigenomic reconstruction as measured by R-squared (b) against different number of hidden nodes with modifications to the LSTM. The y-axis shows the mAP (a) and R-squared (b) and the x-axis shows the number of hidden nodes. The legend refers to the modifications to the Epi-LSTM (shown in blue).

mAP for gene expression of Epi-LSTM with increasing number of hidden nodes (Figure 5a) and saw that bidirectional as well as two-layer LSTMs did not yield significant improvements in downstream performance. On the other hand, using dropout and removing layer norm were detrimental to downstream performance. R-squared for epigenomic reconstruction with the given variations plotted against number of hidden nodes (Figure 5b), reveals a similar trend.

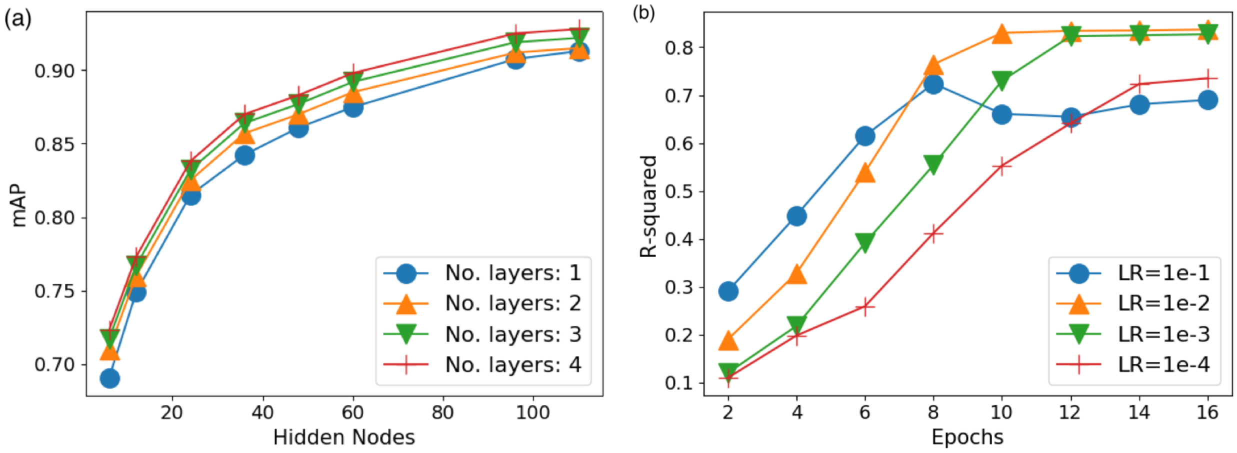


Fig. 6: (a) The average precision for predicting gene expression as measured by mAP for different number of hidden nodes with different number of layers in the LSTM. The y-axis shows the mAP and the x-axis shows the number of hidden nodes. The legend refers to the number of layers used in the LSTM. (b) Regression performance of epigenomic reconstruction as measured by R-squared over increasing number of epochs with different learning rates. The y-axis shows the R-squared and the x-axis shows the number of epochs. The legend refers to the different learning rates.

To verify the marginal increase in mAP with increasing number of layers, we plot the mAP for gene expression with increasing number of nodes using different number of layers (Figure 6a). We see that increasing the number of layers from 1 to 4 yields only minimal increase in mAP and that mAP increases noticeably with increase in number of hidden nodes, making 110 as a satisfactory choice for the hidden size.

In order to chose the optimal value of learning rate to train the Epi-LSTM, we plot the R-squared over epochs with different learning rates (Figure 6b). We observe that while a learning rate of 1e-1 enables faster learning, the R-squared degrades over epochs. Learning rates of 1e-3 and 1e-4 achieve good R-squared but take much longer to learn. A learning rate of 1e-2 seems to find reasonable balance between learning time and R-squared and hence we set our learning rate to this value.

*B. XGBoost*

XGBoost is a gradient boosted random forest. The main parameters of XGBoost deal with booster choice and the parameters for the selected booster. We use the default booster “gbtree” and to make our model slightly less conservative, we chose the default values for regularization parameters such as eta (step size shrinkage), gamma, min_child_weight, max_delta_step, lambda (L2 regularization) and alpha (L1 regularization). We focus on two main parameters pertaining to decision trees that impact expressive power and overfitting, namely: (a) number of estimators: The number of decision trees to be used, and (b) maximum depth: The maximum depth that is allowed for each tree.

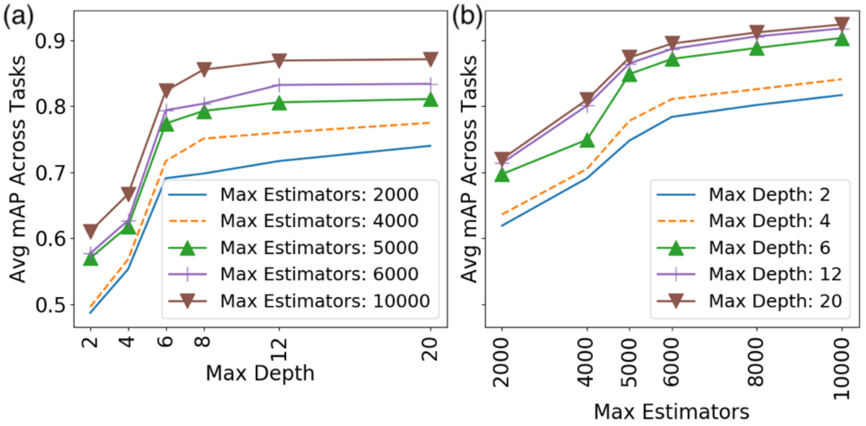


Fig. 7: The average precision for predicting downstream tasks except PE-interactions as measured by mAP. (a) plotted against the maximum depth for different number of estimators. The y-axis shows the mAP and the x-axis shows the maximum depth. The legend shows the different number of estimators. (b) plotted against number of estimators for different maximum depths. The y-axis shows the mAP and the x-axis shows the number of estimators. The legend shows the different maximum depths.

We perform two experiments. In our first experiment, we plot the average mAP for downstream tasks except PE-interactions against the maximum depth for different number of estimators and find the elbow at a maximum depth of 6 (Figure 7a). We also find that 5000 estimators perform comparable to higher number of estimators for the amount of time it takes to run and hence chose the number of estimators to be 5000 (Figure 7a). In our second experiment, we plot the average mAP for downstream tasks except PE-interactions against number of estimators for different maximum depths (Figure 7b) and observe similar trends. We note the elbow at a number of estimators of 5000 and see that a maximum depth of 6 performs close to optimal (Figure 7b).

#### IV. Additional Classification Metrics

Apart from the mean Average Precision (mAP) (which is the same as the Area under PR-curve), we now plot two additional metrics, i.e., AuROC (Area under the ROC-curve), and accuracy.

The Receiver Operating Characteristic (ROC) curve, is a plot of the true positive rate (TPR) versus the false positive rate (FPR) for different classifier thresholds. TPR is calculated by dividing the number of true positives (TP) by the sum of the number of true positives (TP) and the number of false negatives (FN), i.e., $TPR = TP/(TP+FN)$. It is also referred to as the sensitivity. FPR is calculated by dividing the number of false positives (FP) by the sum of the number of false positives (FP) and the number of true negatives (TN), i.e., $FPR = FP/(FP+TN)$. It is also referred to as the inverted specificity. The Area under the ROC-curve (AuROC) reveals how well the model achieves a trade-off between sensitivity and inverted specificity. The higher the AuROC, the higher the model skill.

We evaluate the AuROC for the downstream genomic tasks (averaged across the available cell types) using representations from different methods as input to the XGBoost classifier (Figure 8a) and see that the Epi-LSTM representations have higher AuROC than other methods for all tasks except gene expression, in which, Epi-LSTM and Avocado have similar AuROC (Figure 8a). We also compute the accuracy of classification for the downstream genomic tasks (averaged across the available cell types) using different representations (Figure 8b) and observe a similar trend. The achieved accuracy is higher than the achieved AuROC for all methods across all tasks (Figure 8b).

#### V. Training and Testing Times

In order to check whether the times taken by the Epi-LSTM to perform training and testing are reasonable, we compare our running times with different parameters of the LSTM and with different autoencoders. We consider two important LSTM parameters, namely, the number of hidden layers and the number of hidden nodes.

##### A. LSTM Parameters

We evaluated the training time as a function of the number of hidden nodes for different number of layers (Figure 12a). The time taken is computed for a frame length of 100 inputs on GeForce GTX 1080 Ti GPUs with 16GB RAM and then estimated for a genomic region of 10Mbp. We observe that with a single layer (Epi-LSTM), the amount of time taken to train the model is less than 3.5 seconds per frame; for 4000 frames per 10Mbp, the training takes less than 0.19 days for a 10Mbp region on average (Figure 9a). Furthermore, the training can be significantly sped up by batching multiple inputs and by parallelizing the training across portions of the genome that are sufficiently far apart, rendering

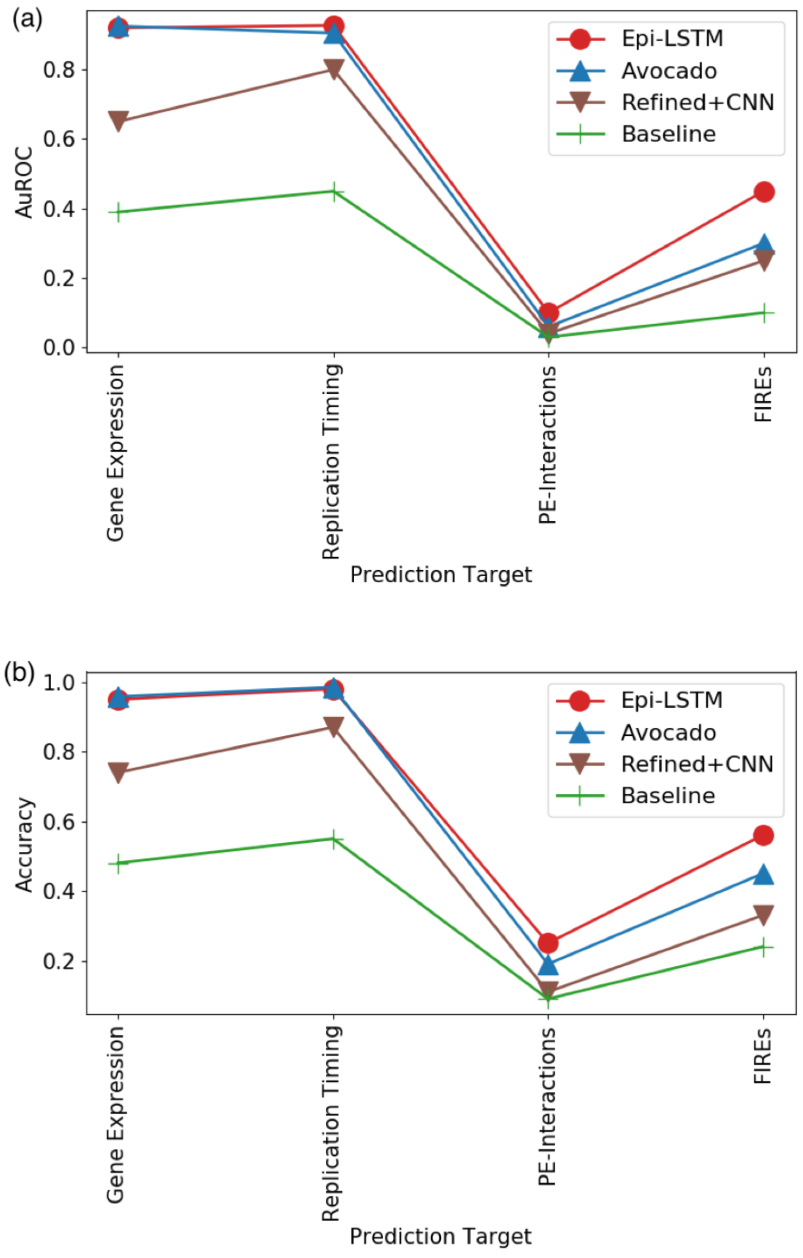


Fig. 8: Prediction performance as measured by AuROC (a) and Accuracy (b) for the downstream genomic tasks (averaged across the available cell types) using representations from different methods as input to the XGBoost classifier. The y-axis shows the AuROC (a) and Accuracy (b) and the x-axis shows the tasks that are prediction targets. The legend shows the different methods.

independence in epigenomic properties. The training time gets worse as you increase the number of layers, increasing about 2 fold for 2 layers and 3 fold for 4 layers (Figure 9a).

In order to evaluate how fast our model can run inference, we evaluated the testing time in seconds (computed per frame length) as a function of the number of hidden nodes for different number of layers (Figure 9b) and observe a similar trend as in training with a order of magnitude reduction. The amount of time taken to run inference is less than 0.7 seconds per frame; for 4000 frames per 10Mbp, the inference takes less than 0.8 hours for a 10Mbp region on average (Figure 9b).

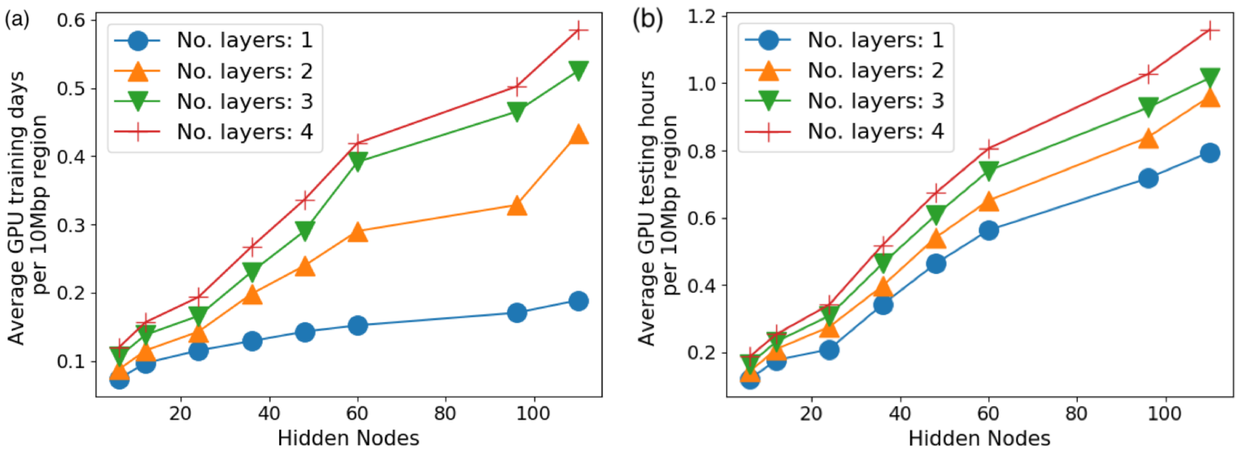


Fig. 9: Training time (a) and Testing Time (b) of Epi-LSTM for different number of hidden nodes with different number of LSTM layers. The y-axis shows the training time in average GPU days (a) and testing time in average GPU hours (b) for 10Mbp and the x-axis shows the number of hidden nodes. The legend shows the different number of LSTM layers. The time taken is computed for a frame length of 100 inputs (0.0183 percent of the chromosome on average) on GeForce GTX 1080 Ti GPUs with 16GB RAM and then estimated for a genomic region of 10Mbp.

##### *B. Other Methods and Autoencoders*

We compare the training and testing time of the Epi-LSTM with other methods like Avocado and REFINED+CNN, as well as architectural variations like RNN, CNN and feed-forward Autoencoders (Table 1). The time taken is computed for a frame length of 100 inputs on GeForce GTX 1080 Ti GPUs with 16GB RAM and then estimated for a genomic region of 10Mbp. The number of hidden layers and hidden nodes for all models are set to be the same.

We see that the recurrent models take slightly longer to train and test when compared to Avocado, convolutional models and feed-forward models (Table 1). The convolutional models take the next highest amount of time followed by Avocado and the feed-forward models (Table 1). The autoencoders can run inference on epigenomic data that the model has not seen before in contrast to Avocado, which has to perform genome-wide training. In this light, the Epi-LSTM is a reasonable choice with respect to training time.

TABLE I: Training and testing time in average GPU days for a 10Mbp region using Epi-LSTM and different autoencoders like LSTM, RNN, CNN and feed-forward autoencoders as well as different methods like Avocado and REFINED+CNN. The time taken is computed for a frame length of 100 inputs (0.0183 percent of the chromosome on average) on GeForce GTX 1080 Ti GPUs with 16GB RAM and then estimated for a genomic region of 10Mbp.

| Model | Training Time (average GPU days per 10Mbp) | Testing Time (average GPU hours per 10Mbp) |
| --- | --- | --- |
| Epi-LSTM | 0.188 | 0.791 |
| Avocado | 0.165 | 0.576 |
| RNN | 0.182 | 0.768 |
| CNN | 0.172 | 0.744 |
| REFINED+CNN | 0.175 | 0.765 |
| Feed-forward | 0.142 | 0.504 |

#### References


[1] The Roadmap Epigenomics Mapping Consortium. [Online]. Available: http://www.roadmapepigenomics.org/

[2] X. J. Mao, C. Shen, & Y. B. Yang. Image restoration using convolutional auto-encoders with symmetric skip connections. arXiv preprint arXiv:1606.08921. 2016.

[3] J. Schreiber, R. Singh, J. Bilmes, & W. S. Noble. A pitfall for machine learning methods aiming to predict across cell types. Genome biology, 21(1), 1-6. 2020.

[4] S. Tuna, & M. Niranjan. Classification with binary gene expressions. Journal of Biomedical Science and Engineering, 2(6), 390-399. 2009.

[5] S. Draghici, P. Khatri, A. C. Eklund, & Z. Szallasi. Reliability and reproducibility issues in DNA mi-croarray measurements. Trends in Genetics, 22(2), 101-109. 2006.

[6] D. Geman, C. d'Avignon, D. Q. Naiman, & R. L. Winslow. Classifying gene expression profiles from pairwise mRNA comparisons. Statistical Applications in Genetics and Molecular Biology, 3. 2004.

[7] S. Tuna, & M. Niranjan. Inference from low precision transcriptome data representation. Journal of Signal Processing Systems, 58(3), 267-279. 2009.

[8] N. Friedman, M. Linial, I. Nachman, & D. Pe'er. Using Bayesian networks to analyze expression data. Journal of Computer Biology, 7(3–4), 601–620. 2000.

[9] D. Pe'er, A. Regev, G. Elidan, & N. Friedman. Inferring subnetworks from perturbed expression profiles. Bioinformatics, 17(1), S215–224. 2001.

[10] X. Zhou, X. Wang, & E. R. Dougherty. Binarization of microarray data on the basis of a mixturemodel. Molecular Cancer Therapeutics, 2(7), 679–684. 2003.

[11] I. Shmulevich, & W. Zhang. Binary analysis and optimization-based normalization of gene expression data. Bioinformatics, 18(4), 555–565, 2002.

[12] H. I. H. Chen, Y. C. Chiu, T.Zhang, S. Zhang, Y. Huang, & Y. Chen. GSAE: an autoencoder with embedded gene-set nodes for genomics functional characterization. BMC systems biology, 12(8), 45-57. 2018.

[13] O. Bazgir, R. Zhang, S. R. Dhruba, R. Rahman, S. Ghosh, & R. Pal. Representation of features as images with neighborhood dependencies for compatibility with convolutional neural networks. Nature communications, 11(1), 1-13. 2020.
